# Multi-Objective Approach for Identifying Cancer Subnetwork Markers

**DOI:** 10.1101/2020.09.18.304089

**Authors:** Faezeh Bayat, Mansoor Davoodi

## Abstract

Identifying genetic markers for cancer is one of the main challenges in the recent researches. Between different cohorts of genetic markers such as genes or a group of genes like pathways or sub-network, identifying functional modules like subnetwork markers has been more challenging. Network-based classification methods have been successfully used for finding effective cancer subnetwork markers. Combination of metabolic networks and molecular profiles of tumor samples has led researchers to a more accurate prediction of subnetwork markers. However, topological features of the network have not been considered in the activity of the subnetwork. Here, we apply a novel protein-protein interaction network-based classification method that considers topological features of the network in addition to the expression profiles of the samples. We have considered the problem of identifying cancer subnetwork markers as a multi-objective problem which in this approach, each subnetwork’s activity level is measured according to both objectives of the problem; Differential expression level of the genes and topological features of the nodes in the network. We found that the subnetwork markers identified by this method achieve higher performance in the classification of cancer outcome in comparison to the other subnetwork markers.

## 1 Introduction

Distinguishing between different types of tumors is a fundamental issue in diagnosis of cancer. While outward characteristic of some tumors might be similar, they may not react to the treatment in the same way [18]. It may be related to the corresponding underlying mechanism of the disease-causing genes. Hence, it might be more beneficial if treatments become more personalized according to each type of tumor with certain characteristics [20]. Therefore, in order to identify the underlying mechanism of cancerous cells, cancer markers were introduced [8]. With the development of the high-throughput gene expression data, identifying the biomarkers has been more accurate [18].

Identifying the markers of diseases, reveals new insights in studying the biological underlying mechanism of cells. Cancer has become one of the death-causing diseases which leads the researchers to find the predictive biomarkers which are responsible for cancer clinical outcome or treatment response [21, 2, 9]. Earlier studies introduced predictive biomarkers as single genes which despite some drawbacks, have shown significant results. Gene markers consider the most differentially expressed genes as markers [19, 13]. In these approaches, functional relation between genes has not been considered. With the advent of interaction networks such as protein-protein interaction networks, which show the physical or functional relation between genes or gene products, it has been possible to combine the gene expression data with the interaction networks in order to consider the functional relation between genes.

In order to overcome gene marker’s defects, new aspects of biomarkers were introduced; Metagene markers [16, 1]. Metagene markers which include gene sets, pathways and subnetworks are more robust solutions for biomarkers. The idea behind the metagene markers is the fundamentally modular underlying mechanism of cellular organizations. In other words, genes’ physical or functional interactions have significant effect on the behavior of the cell. Functional modules which consist of active modules, conserved modules, differential network modules and composite functional modules are the group of genes, gene products or gene sets that are in connection with each other or they are functionally coordinated [14]. A pathway or subnetwork refers to a connected region in a network which has significant changes in expression of its consisting genes in different conditions [14].

Identifying functional modules through integrating high-throughput data such as protein-protein networks, metabolic networks and gene expression data has shown more reliable results [12]. In order to find functional modules such as pathways or subnetworks, activity of each module should be measured. To this end, weight functions should be assigned for each of the modules.

There are different approaches for assigning a score for modules. Gene-based methods, interaction-base methods, group-based methods and probabilistic models are some of the methods that are used for scoring modules [5]. Between all the subnetworks, the ones with the highest weight are the active subnetworks. It has been proved that finding the connected subnetwork with maximal weight is an NP-hard problem [8]. One of the earliest approaches for finding active pathways was introduced by Ideker et al [8]. In order to identify the active pathways, activity of a subnetwork is derived from a gene-based scoring method. Each gene in the network is assigned a z-score based on its differential expression in the gene expression data and the activity level of the subnetwork is the aggregation of its consisting gene’s activity. For finding the highest weight subnetwork, simulated annealing algorithm is applied on the network. Another approach that was introduced by Guo et al. used interaction-based method for scoring the subnetworks [7]. For this purpose, a score is assigned to each interaction between proteins and the overall activity of a subnetwork is the aggregation of the interactions’ scores in the subnetwork. Nacu et al. and Segal et al. used a group-based and probabilistic scoring method respectively [15, 17]. One of the most recent methods has modeled the problem of finding cancer subnetwork markers as a game theoretic approach which considers the proteins in the network as players of the game [6]. The purpose of the game is to select the optimal set of players which maximize the final scoring function that they have defined for each player.

In the aforementioned scoring methods, topological features and structure of a network were not considered to our knowledge. Therefore, we propose a new method for assigning a weight to subnetworks which considers different topological features of the subnetwork in finding the most discriminative subnetworks in addition to the differential expression of its consisting genes. In order to evaluate the efficiency of the proposed method, we compared it with the state-of-the-art studies. We compared the results of the proposed multi-objective approach with game theoretic approach [6], optimally discriminative [4], greedy method [3], pathway-based and gene-based methods. As results show, proposed method achieve higher classification performance in cancer outcome in comparison with the other approaches.

## 2 Material and Methods

### 2.1 RNA-seq data

In order to evaluate the efficiency of the proposed multi-objective method, we used the data presented in one of the recently published papers [6]. Three independent datasets for gene expression profiles were used. Two of these datasets are available with GEO accession numbers GSE7390 for Belgium dataset and GSE1456 for the Sweden dataset. The last dataset (Netherland dataset) is taken from van de Vijver et al. [19]. Belgium and Sweden datasets are profiled on the Affymetrix U133A platform and have 198 and 159 samples respectively. The samples are divided into metastatic and non-metastatic labels and both of these datasets have 35 metastatic samples. Netherland dataset is profiled on the Agilent microarray platform and have 295 samples which 78 of them are metastatic.

### 2.2 Protein-protein interaction networks

For the PPI network, human PPIN created by Lage et al. is used [11]. PPI network contains 12894 nodes and 426579 interactions. For the integration of gene expression data and PPI network, the microarray data should overlay on the PPI network. For this purpose, each gene in the gene expression data should be mapped to its corresponding protein in the PPI network and if there is no mapping for the protein, it should be removed from the network.

### 2.3 Multi-Objective approach

As mentioned before, purpose of this paper is to find the closely connected subnetworks which are more influential than the others in causing cancer. In other words, these subnetworks must have the most discriminative power in separating cancerous samples from non-cancerous ones. In order to distinguish between different subnetworks, we used different objectives for each subnetwork. The first objective that is very important in causing a cell in becoming cancerous, is the different expression level of the genes. In other words, down-regulated or upregulated genes tend to participate more in manipulating cells toward becoming cancerous. Therefore, subnetworks that most of its genes have differentially expressed genes are more probable to be one of the discriminative subnetworks. The second objective that is taken by multi-objective approach is to consider the topological features of the network. Beside the differential expression of the genes which plays a critical role in making a cell cancerous, it seems that some proteins with specific characteristics have the potential to speed up the process of making a cell cancerous. For example, suppose that most of the genes are effected due to their down/up-regulation. Here, due to the high degree of the hub proteins, it is more probable that these proteins play a major role in spreading the disease. In this paper, we considered some centrality measurements and coefficients that should satisfy the multi-objective criteria.

### 2.4 Feature selection

In order to define the objectives of the problem, we need to explain what features are used in this approach.

#### 2.4.1 First objective: Differential expression level of the genes

In order to specify the genes with the differential expression, some metrics have been introduced, e.g. z-score and t-score. Here, we used t-score metric and a metric that was introduced in Optimally discriminative approach [4] as a distance-based function which is shown in the following equation:

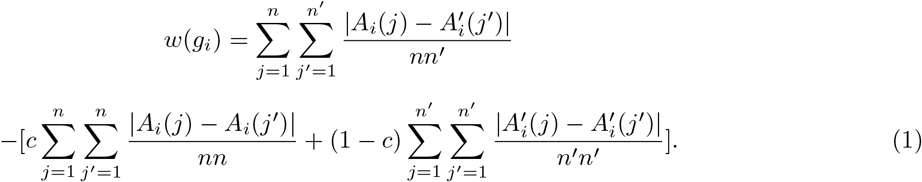

where *w*(*g*(*i*)) shows the weight of each gene. Suppose that we show the expression matrices of the positive and negative samples by *A* and *A′* and notations *n* and *n′* shows the number of samples in positive and negative classes respectively. 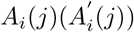 denotes the expression of gene *g*(*i*) in sample j in matrix *A*(*A′*). The idea behind the distance-based function is to define the differential expression level of the genes as the difference between the total distance of samples from all classes and the total distance between the samples from the same class [4]. Therefore, weight of a subnetwork S consisting of k genes is measured by the following equation:

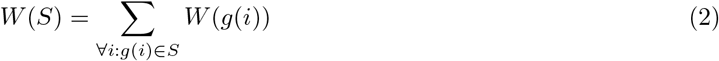

Similarly, weight of a subnetwork regarding to its consisting genes’ t-score can be calculated with the aforementioned equation which *w*(*g*(*i*)) shows gene *g_i_* score gained by the t-test.

#### 2.4.2 Second objective: Topological features of the nodes

As it is mentioned in the previous sections, some nodes in the network with specific characteristics can amplify the effects of the differentially expressed genes which are more probable to be disease causing genes. Therefore, in this study, we are going to investigate the impact of some of the most important features of the nodes in the network in their ability to distinguish different samples from each other in addition to the differential expression level of the genes. For this purpose, we considered following centralities and coefficients for the nodes of the network. Betweenness centrality, bridging centrality, brokering coefficient, clustering coefficient, degree centrality and local average centrality are the metrics that we used as topological features of the nodes in the network [10].

### 2.5 Mathematical modeling of the subnetworks

In order to find the connected subgraphs, we considered subnetworks as the shortest paths between the nodes representing proteins. To show a discriminative power of a subnetwork, we need to assign a weight to the subnetwork as a metric for showing its activity level. Suppose that shortest path between two nodes which is representing a subnetwork is shown by *A* which has *k* genes. According to the multi-objective approach, each subnetwork should satisfy the objectives of the problem; the first objective is to maximize the overall activity of the subnetwork according to its differentially expressed genes which are more likely to be the intended genes. To this purpose, we assign a weight function to each subnetwork as the sum of its genes’ differential expression scores which are gain from one of the two metrics that were introduced before, t-score or distance-based function. Therefore, we consider the weight of the subnetwork *A* as:

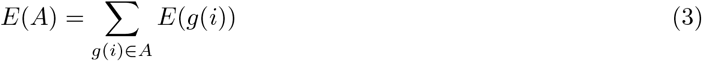

where *E*(*g*(*i*)) shows the differential expression score of the gene *g*(*i*) in the subnetwork. To sum up, for this step of the approach, the subnetworks with the highest weights are the candidates for becoming cancer subnetwork markers. In order to evaluate the subnetworks according to their topological features, again we need to assign a weight to each of them. Therefore, a weight function is defined as follows:

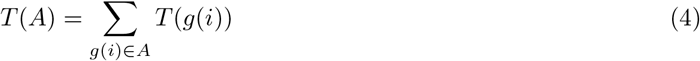

where *T* (*g*(*i*)) shows one of the coefficients or centrality measures of the gene *g*(*i*) in the subnetwork. Now, each subnetwork is shown with two objectives like Fig.1.

**Fig. 1.**
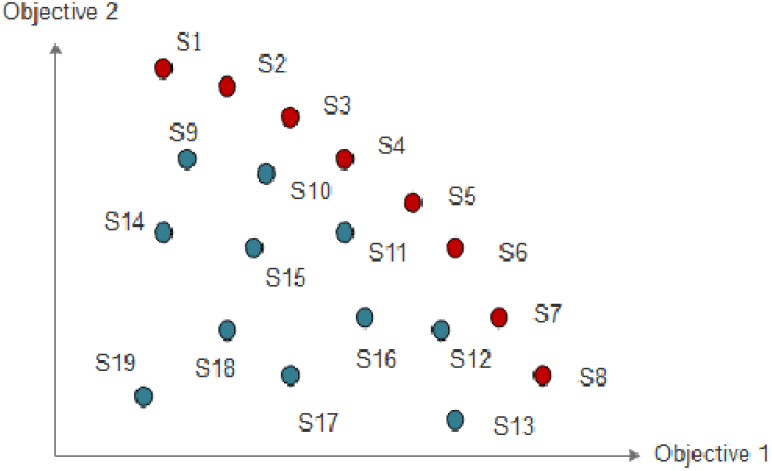
Each subnetwork is shown by its two objectives

According to the multi-objective approach, in order to identify the subnetworks which can satisfy the both objectives, we need to find the Pareto optimal front of the subnetworks which are shown in red dots. Pareto optimization is used for optimizing problems with multiple criteria. In other words, Pareto optimal seeks to optimize the objectives of the problem simultaneously. Therefore, subnetworks in a Pareto optimal front are the optimal solutions for the problem of finding the most discriminative subnetworks. Due to the complexity of finding the Pareto optimal front of the two-objective problem, we considered the problem as a one-objective problem. For this purpose, we combined the two objectives of the problem in one weight function as follows:

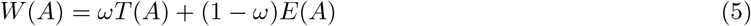

where *ω* varies in the range of [0, 1]. By considering enough samples of the weight function for different *ω* levels, we make sure that the large portion of the optimal solution are included. To this end, we considered *ω* starting from 0 and each time increased by 0.1 till 1. In the final step, we introduce *N* optimal subnetworks which in the previous works, *N* is set to 50. Therefore, among all the subnetworks for each *ω*, we select the 50 top subnetworks according to their weight. In other words, 50 subnetworks with the highest weights are selected as the optimal solution of the problem. Pseudo code of the proposed method is provided in the following.

#### Algorithm 1

Multi-objective approach

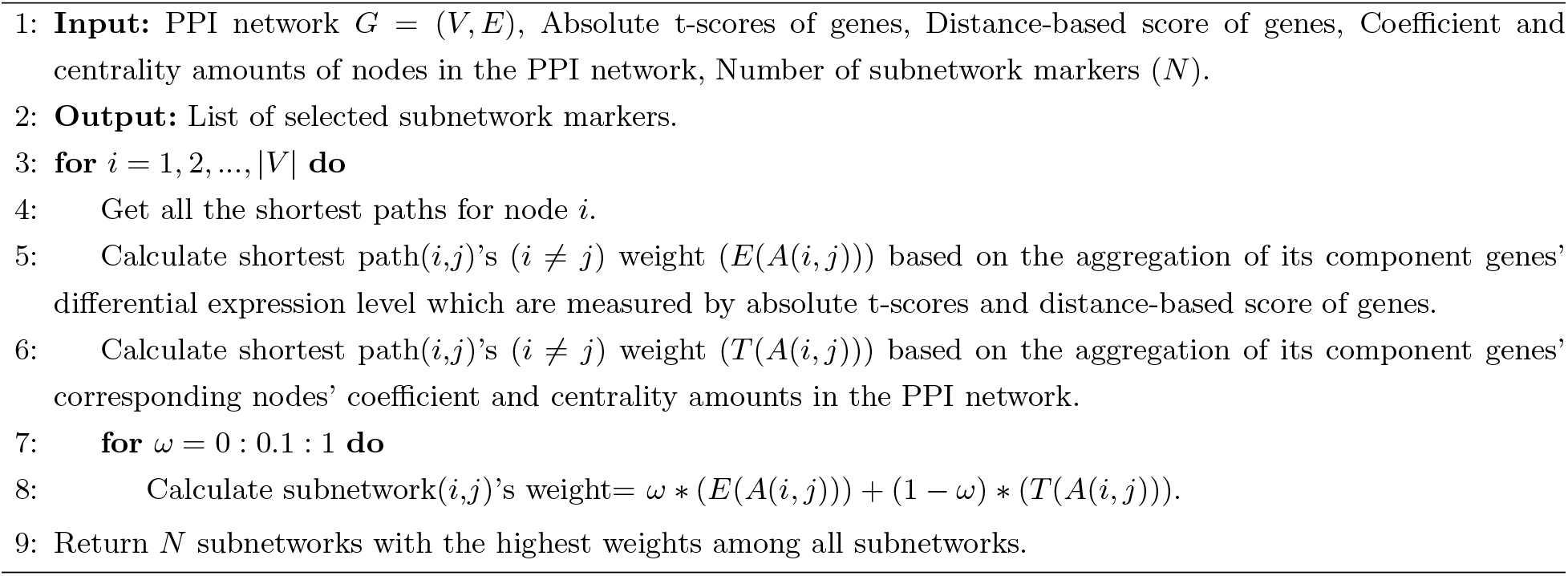

## 3 Results and discussion

In order to identify subnetwork markers, different criteria are taken into account. First set of these criteria considers topological features of the proteins in the network. Second set is concerned with the differential expression level of the genes in the microarray dataset. Here, we considered six topological features of the nodes and two different functions for measuring differential expression of the genes. Therefore, as it is mentioned in the methods section, twelve sets of subnetwork markers are identified by our multi-objective approach. Thus, for evaluation of these subnetwork markers, classification performance and gene ontology analysis have been considered.

In order to evaluate the performance of the proposed method, we used support vector machine (SVM) method for within-dataset and cross-dataset analysis. We compared our results with different approaches for identifying cancer markers. Single gene approach, pathway-based and network-based approaches are the methods that we compared our method to. In each of these approaches, the top 50 markers are extracted and introduced as the cancer markers. For evaluation of the network-based methods, 50 top subnetwork markers are used to train and test the SVM classifier. For both within-dataset and cross-dataset analysis, we repeated five-fold cross validation ten times. For comparing the results, classification accuracy (ACC) and area under curve (AUC) criteria are chosen.

### 3.1 Considering topological features of the PPI networks alongside the differential expression of the genes outperform the classification performance

For evaluating the classification performance, we first analyzed the within-dataset results. In this procedure, subnetwork marker identification approaches were applied on each dataset and top subnetwork markers were selected. These markeres were then used to train and test the SVM classifier. Results of the within-dataset analysis is shown in the Figure 2a. As it is shown, it seems that pair of objectives where *tscore* is considered as the differential expression function, have better performance in comparison with the *distance base* function. Among the different topological features of the nodes in the *PPI* network, *Betweenness centrality* has shown better classification performance in the within-dataset analysis. The results provided in Figure 2a is averaged between three *Netherland*, *Sweden* and *Belgium* datasets. More detailed classification performance results are provided in the Supplementary Information Table 2.

**Fig. 2.**
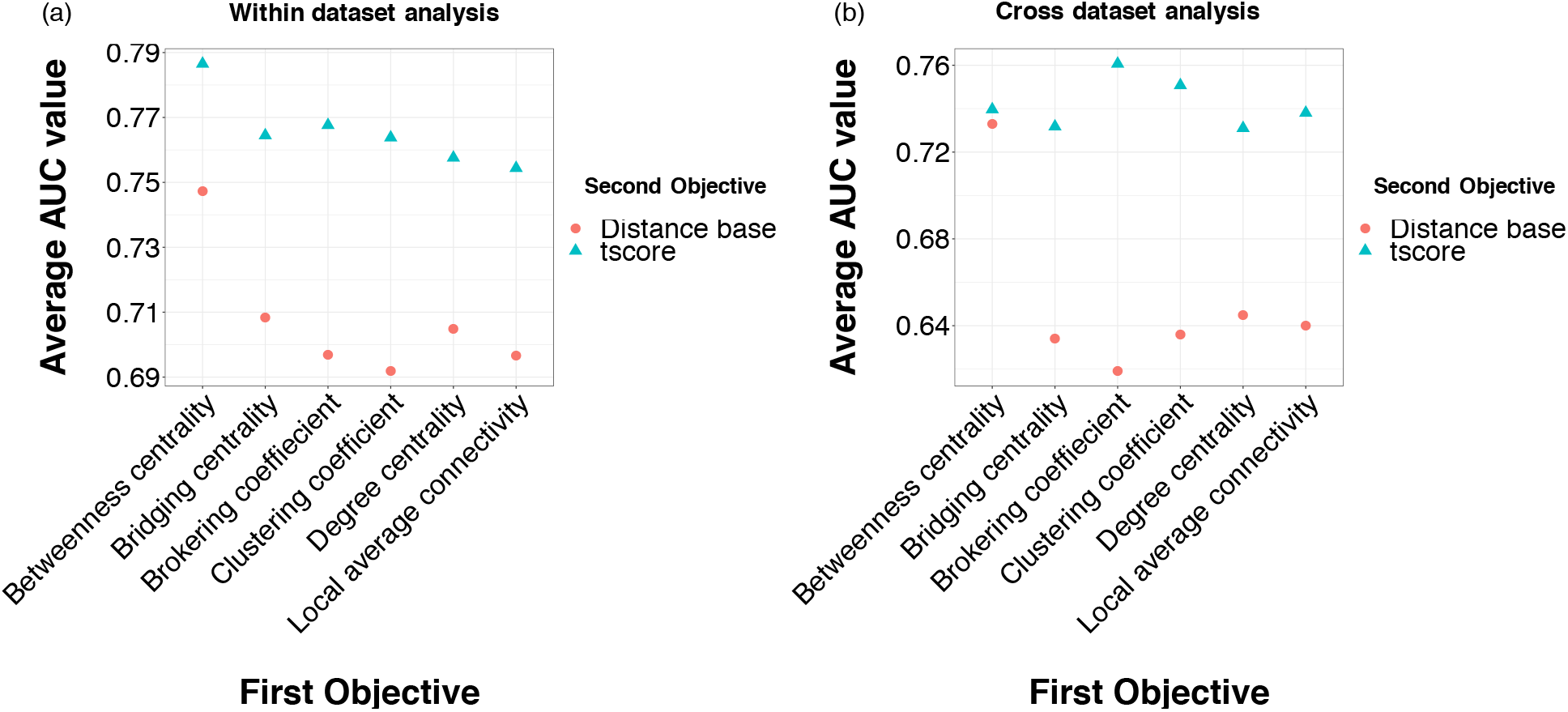
Multi-objective classification performance. a) average performance across all datasets in within-datset analysis. b) average performance across all three sets of experiments where subnetworks are extracted from one dataset and testes on the other two datasets in cross-dataset analysis.

Among the twelve different pairs of objectives in the multi-objective approach, we selected *Betweenness centrality-tscore* as the candidate pair to compare our resutls to the other well known subnetwork marker identification approaches. The results are shown in the Figure 3. As it is shown, multi-objective approach is performing approximately similar to the GTA approach and both of these methods outperform the other four classification methods including SGM, OptDis, Pathway-based and Greedy method in both two classification evaluation metrics; ACC, accuracy and AUC, area under the curve (also known as AUROC, Area Under the Receiver Operating Characteristics).

**Fig. 3.**
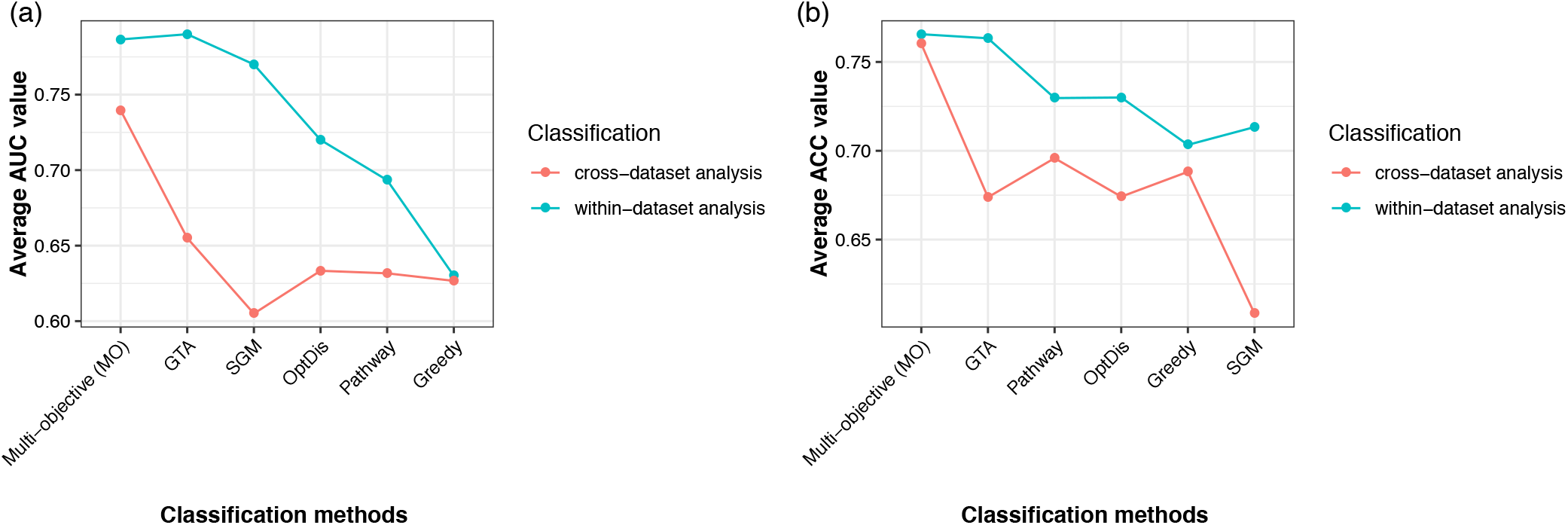
Classification performance comparison between all approaches a) AUC metric b) ACC metric in within-dataset and cross-dataset analysis.

### 3.2 Are the identified subnetworks reproducible?

In order to verify the reproducibility of the selected subnetwork markers, cross-dataset analysis was performed. For this purpose, selected subnetwork markers, identified from each dataset, were used to train the SVM classifier. Then the trained classifier were tested on the other two datsets. As mentioned before, five-fold cross-validation is repeated ten times. The cross-dataset validation results for multi-objective approach are shown in Figure 2b. Similar to the within-dataset analysis results, it seems that *tscore* function for the differential expression is performing better than the *distance-base* function. Also, *Brokering coefficient* shows better classification performance in combination with the tscore function.

The results provided in Figure 2b are averaged between three sets of experiments where in each set, sub-network markers from one dataset have been used for the reproducibility purpose on the other two datasets. More detailed classification performance results for cross-dataset analysis are provided in the Supplementary Information Table 3-5.

Following the within-dataset analysis, we selected *Betweenness centrality-tscore* as the candidate pair to compare our resutls to the other approaches. Eventhough *Brokering coefficient-tscore* is performing better in cross-dataset analysis, we needed to choose same set of objective pairs as in within-datset analysis to have a fair comparison. The results are shown in the Figure 3. As it is shown, multi-objective approach is outperforming all the other approaches in terms of ACC and AUC. This implies that generalizability based on the multi-objective approach is one of its key advantages.

In addition to the classification performance analysis, we analyzed the selected subnetworks from three datasets in order to find the overlapping genes among these datasets. For computing the overlapping amount for each pair of datasets, intersection of the two unique gene sets is divided by the sum of the two unique gene sets. The results are shown and compared to the other approaches in Table 1.

**Table 1.** Overlap of the selected subnetwork markers in different methods.

| Methods | Belgium-Sweden | Belgium-Netherlands | Netherlands-Sweden |
| --- | --- | --- | --- |
| MO:Betweenness centrality-tscore(%) | 100 | 18.42 | 18.42 |
| MO:Bridging centrality-tscore(%) | 25 | 2.08 | 1 |
| MO:Brokering coefficient-tscore(%) | 25.58 | 2.08 | 1.16 |
| MO:Clustering coefficient-tscore(%) | 24.21 | 4 | 1.05 |
| MO:Degree centrality-tscore(%) | 97.72 | 13.95 | 16.27 |
| MO:Local average connectivity-tscore(%) | 86.84 | 47.36 | 40.47 |
| MO:Betweenness centrality-Distance-base(%) | 100 | 18.42 | 18.42 |
| MO:Bridging centrality-Distance-base(%) | 33.33 | 0 | 1 |
| MO:Brokering coefficient-Distance-base(%) | 35 | 1.75 | 1.35 |
| MO:Clustering coefficient-Distance-base(%) | 87.5 | 38.19 | 37.33 |
| MO:Degree centrality-Distance-base(%) | 100 | 14.63 | 14.63 |
| MO:Local average connectivity-Distance-base(%) | 100 | 51 | 51.42 |
| GTA(%) | 16.24 | 11.44 | 13.80 |
| Greedy method(%) | 11.57 | 8.84 | 7.92 |
| OptDis(%) | 9.69 | 10.53 | 12.48 |

### 3.3 Single-objective or multi-objective?

Furthermore, In order to determine the impact of each objective of the problem in classification performance, we compared the results of the multi-objective approach in classification of the samples with the single-objective approach. Therefore, we considered the problem of identifying subnetwork markers in two conditions; Multi-objective approach and single-objective approach. In the single-objective mode, once we considered the topological features of the problem as the only objective of the problem and once we considered the differential expression of genes as the objective of the problem. Then we compared the impact of each objective in the classification performance in within-dataset analysis. The results of the multi-objective and single-objective approach are shown in Figure 4a and Figure 4b respectively. We compared the multi-objective approach with single-objective method by getting the average over the twelve possible combinations of the objectives and as results in Figure 4a shows, multi-objective approach outperforms the single-objective method.

**Fig. 4.**
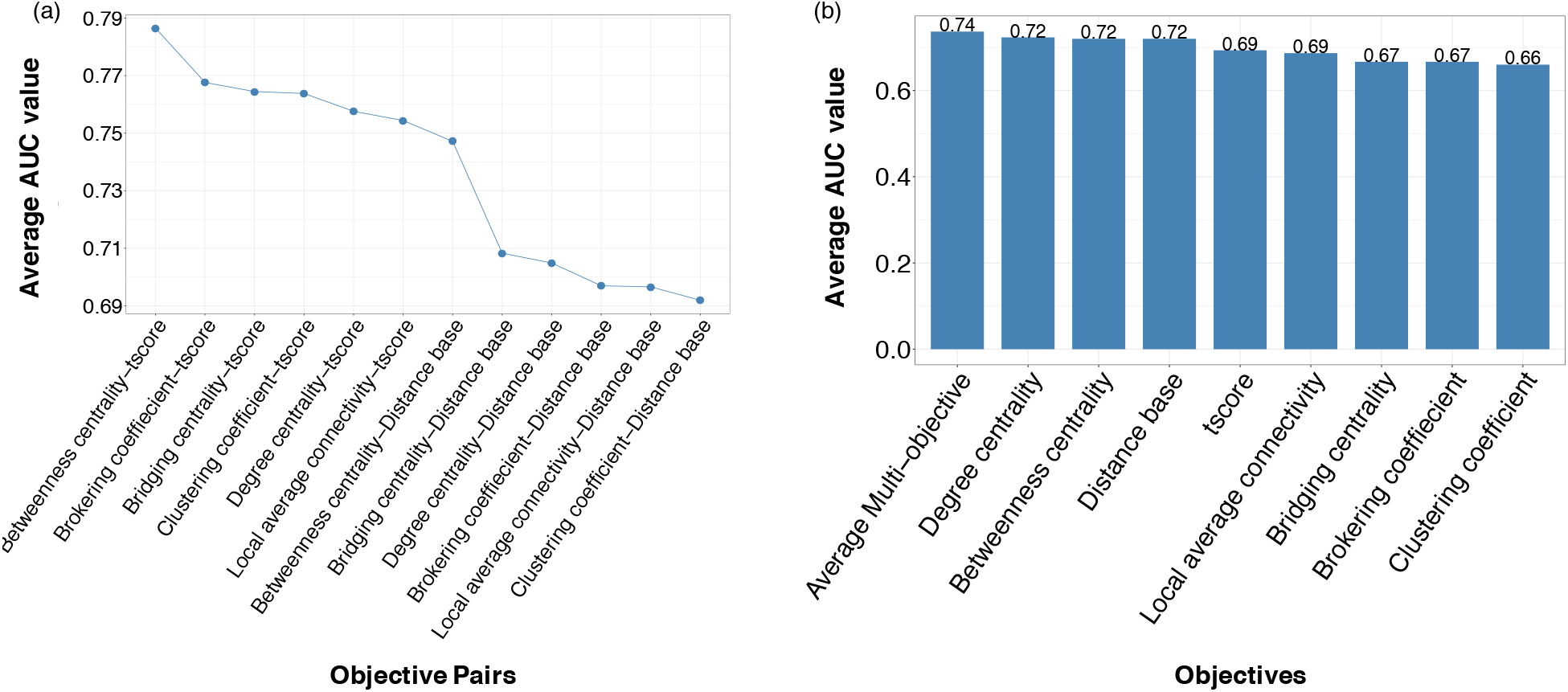
Comparison of the multi-objective and single-objective approaches a) multi-objective b) single-objective (The first column is numerical average of the multi-objective approaches in (a)).

### 3.4 Enrichment analysis

In order to analyze enriched biological processes of the identified subnetwork markers, gene ontology (GO) analysis is performed. For this purpose, we considered the different multi-objective methods separately in order to demonstrate the impact of each pair of objectives on gene ontology analysis. Therefore, we divided the analysis into six groups. Each group contains the subnetworks with one topological feature at a time and for the second objective, both distance-base function and t-score were considered. Following the classification performance analysis, here we just demonstrate the results of the *Betweenness centrality* enrichment analysis over all three datasets which is shown in Figure 5. The results of the remaining topological fetures can be found in the Supplementary Information Figures 6-10. As results in Figure 5 shows, almost all the subnetworks which their topological feature of their network is *betweenness centrality* have the same enriched biological processes and this is due to the large amount of betweenness centrality of some nodes which overcome the amount of t-score value or distance function value of a subnetwork which results in stucking in a specific subnetworks which just their betweenness centrality value are high. Therefore, because of the same topological structure of PPI for both Belgium and Sweden datasets, identified subnetworks are same for both t-score and distance base function objectives. For Netherland dataset, some differences have been seen between the t-score and distance function. Among all topological features of the nodes in GO analysis, bridging centrality and brokering coefficient have shown superiority in enrichment biological processes such as cell cycle, DNA repair, DNA replication and DNA metabolic process in comparison with the other features. Researchers believe that nodes with higher brokering coefficient tend to be disease-causing genes more than other genes. Brokering coefficient shows the difference between the log-transformed degree of a node and the clustering coefficient. In other words, nodes which have higher degrees and lower clustering coefficients are more probable to be disease causing genes.

**Fig. 5.**
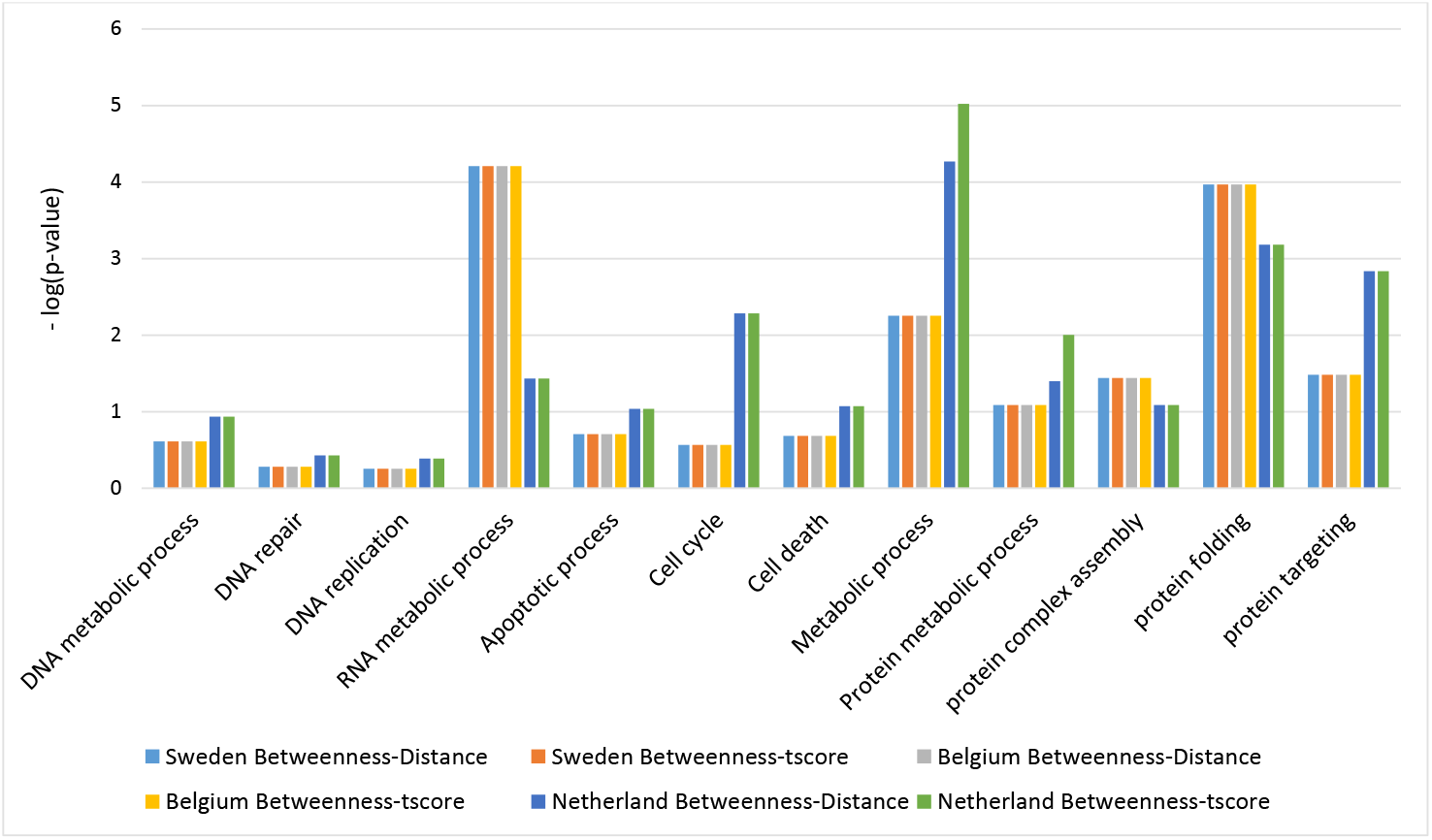
Enriched biological processes of the identified subnetwork markers for Sweden, Netherland and Belgium datasets which the topological feature of the nodes is the betweenness centrality.

## 4 Conclusion

In this paper, we proposed a novel method for identifying cancer subnetwork markers. Most of the previous studies considered differential expression level of the genes as an important factor in defining the activity of the subnetworks; while topological features of the networks were not considered in defining weight function for the subnetworks. The idea behind this study, is to find out the impact of the topological features of the nodes in their discriminative power between different samples in addition to the differential expression level of genes. For this purpose, we considered the problem as a multi-objective problem which one of the objectives is the differential expression of the genes and the other one is the topological features of the nodes. As the results show, this approach overcomes the other subnetwork marker approaches in cancer outcome classification. We considered subnetworks as the shortest path between nodes while if nearly all possible subnetworks with different structures were considered, it was more probable that the right subnetworks were chosen and the results may become better.

